# SciBet: a portable and fast single cell type identifier

**DOI:** 10.1101/645358

**Authors:** Chenwei Li, Baolin Liu, Boxi Kang, Zedao Liu, Yedan Liu, Changya Chen, Xianwen Ren, Zemin Zhang

## Abstract

Fast, robust and technology-independent computational methods are needed for supervised cell type annotation of single-cell RNA sequencing data. We present SciBet, a Bayesian classifier that accurately predicts cell identity for newly sequenced cells or cell clusters. We enable web client deployment of SciBet for rapid local computation without uploading local data to the server. This user-friendly and cross-platform tool can be widely useful for single cell type identification.

## Main

As single cell RNA sequencing (scRNA-seq) datasets grow in number and size, supervised projection of newly-generated data onto annotated labels has become more efficient and practical than unsupervised approaches. One such major challenge is the reliable and rapid cell type identification given a newly sequenced cell. Traditional classification methods such as random forest classifier (RF) and support vector machine (SVM) are often time-consuming, whereas tools specifically designed for such projection trade accuracy for speed^1^. Integration-oriented tools or deep-learning based tools rely on computation-intensive search of anchor cells or iteration of multiple epochs. Such practice can become inefficient if a huge data set like the Human Cell Atlas (HCA)^3^ is used as reference.

Here, we quantify the variability of gene expression in scRNA-seq data with the statistic *entropy*, which can be generalized across datasets, faithfully reflects the variability of gene expression within a population, and allows the direct quantification of its heterogeneity (Online Methods). Based on this statistic, we devised a novel E(ntropy)-test for robust supervised identification of highly informative genes and applied this new test and its basic concept for SciBet (Single cell identifier based on E-test), a Bayesian classifier for scRNA-seq data projection. Using a wide range of scRNA-seq datasets, including data from different biological systems and technologies, we demonstrate that SciBet provides fast, robust, scalable, and accurate classification of cells.

In order to accelerate the process of training and testing of the classifier and identify biologically-informative genes, we first select top differentially-expressed genes across the given cell groups in a supervised manner by E-test. We use the statistic entropy to measure the dispersion degree of the Poission-Gamma mixture distribution of the gene expression in a parametric manner, where the entropy can be directly calculated with the mean gene expression. Based on the clustering result, we use the total difference of entropy among all groups to measure the overall heterogeneity of the gene expression (Fig. 1a and Online Methods). To illustrate the performance and the scalability of E-test, we benchmarked our method against F-test (one-way ANOVA) and M3Drop^12^ using 14 published datasets encompassing data derived from different technologies (Supplementary Table 1). E-test consistently achieved the highest classification accuracy in the cross-validation task within each dataset (Fig. 1b and Supplementary Fig. 1). The superiority of E-test is independent on the number of selected genes and classification algorithms, indicating the robustness of E-test for identifying biologically discriminating genes. We further used nine immune cell types from 7 studies to test the performance of E-test for supervised gene selection (Supplementary Table 2). Notably the top 54 genes with maximal Δ*S* were all well-established immune cell markers, known to play pivotal roles in corresponding cell types^4^ (Fig. 1c). The identified marker genes also allowed better visualization, with distinct immune cell populations across different studies separately located in the 2D UMAP plot^5^, further supporting their biological relevance (Fig. 1d). Since E-test is parameter-free, an extremely fast speed is delivered compared to those traditional gene selection methods. In principle, the computational time of E-test increases linearly with the numbers of cell types, cells and genes, in contrast to the quadratic increase for most other available methods. The time consumption and cell number relationship (Fig. 1e) confirmed this trend, demonstrating the scalability of E-test for extremely large datasets.

**Fig 1.**
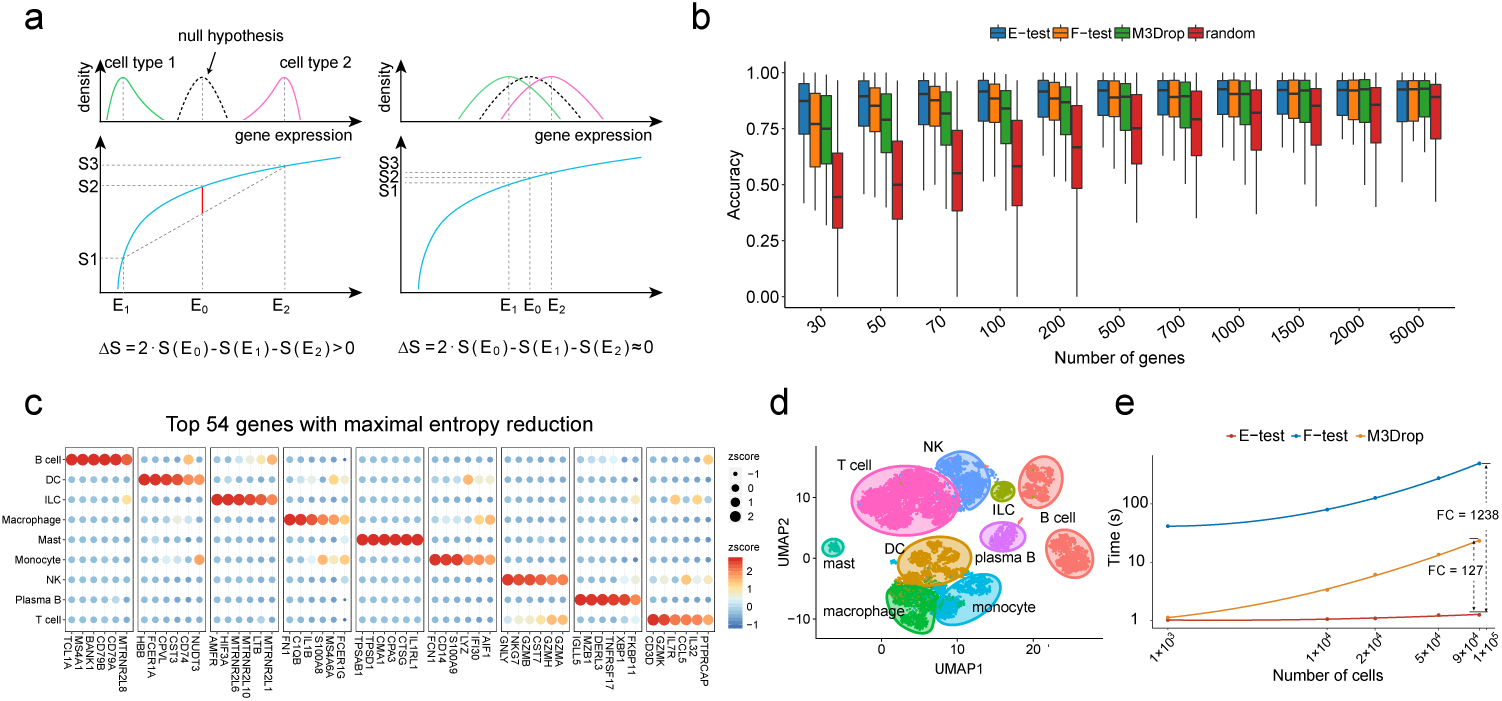
E-test for supervised feature selection. **(a)** Schematic illustration of E-test. Genes with significant differences among cell clusters have large Δ*S* values (left) while Δ*S* of genes with no significant differences tend to be zero-valued (right). Δ*S* can be efficiently calculated by the analytical *S* - *E* function. **(b)** E-test performance is measured by percentage of correctly mapped cells using scmap, with 50-time cross-validations of 14 datasets (Supplementary Table 1). **(c)** Expression heatmap of the top 54 genes selected by E-test for the integrated immune dataset (Supplementary Table 2). **(d)** 2D-UMAP showing the dimensional reduction result based on the genes in **(c). (e)** Single CPU consuming times for gene selection process with E-test, F-test and M3Drop. Solid lines are loess regression fitting (span = 2), implemented with R function geom_smooth.

With the rapid development and application of scRNA-seq, the scale of scRNA-seq dataset has become increasingly large, with the current dataset reaching millions of cells. Accurate and quick determination of single cell identities has become an urgent task. Taking advantages of the remarkable ability of E-test to identify highly informative genes and the intuitive concept that the expression of each cluster is represented by the mean expression vector, we developed SciBet (Single Cell Identifier Based on Entropy Test) for accurate, fast, and robust single cell identification. SciBet determines the identity of each query cell by using a multinomial-distribution model (Fig. 2a, Online Methods) and gives the probabilities of test cells belonging to each of the existing cell types of a reference based on Bayesian decision. It achieves high accuracy and low false positive rate by setting a null dataset as the alternate reference for cells with types not yet covered by the existing data (Online Methods).

**Fig 2.**
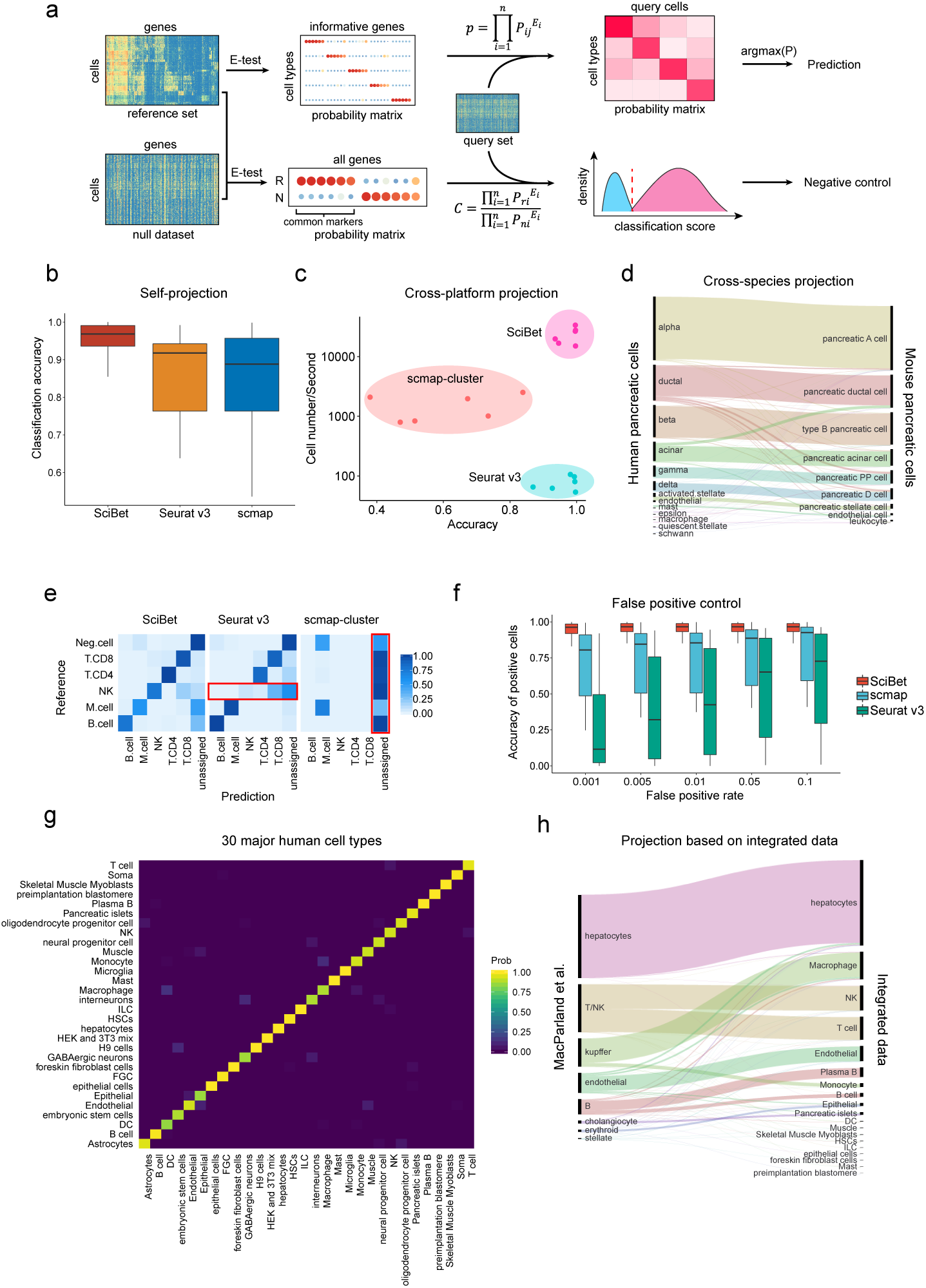
SciBet use and performance. **(a)** Workflow of SciBet. Based on informative genes identified by E-test, each cell type is represented by a probability vector, where each probability value is derived by normalizing mean gene expression. The identification of each query cell is determined using a multinomial-distribution model (Methods). For false positive control, we identify the common markers of reference set using E-test (with a null dataset served as background) and calculate the classification score for each query cell. Query cells with low classification scores are marked as “unassigned”. **(b)** Classification accuracy of self-projection tasks, estimated by SciBet, Seurat v3 and scmap for n = 8 datasets listed in Supplementary Table 3. **(c)** Speed and classification accuracy of cross-platform experiments estimated by SciBet, Seurat v3 and scmap with 6 instances (Supplementary Table 4). **(d)** Cross-species classification with three human pancreas datasets^6–8^ projected to *Tabula Muris* dataset^9^ (Sankey diagram). The height of each linkage line reflects the number of cells. **(e)** Confusion matrix for hold-out evaluation where “Neg.cell” (malignant cells, CAF and endothelial cells) were removed from the reference. Query cells with lowest classification or similarity scores were labeled as “unassigned” (n = 2228, the total number of “Neg.cell”). **(f)** False positive evaluation with cell types not present in reference as negative control, estimated by SciBet, Seurat v3 and scmap with n = 8 cases (Supplementary Table 5). **(g)** Confusion matrix of cross-validation evaluation for 30 integrated major human cell types (Supplementary Table 1-5). (h) Single cell classification for a human liver dataset^11^ with integrated human dataset as reference, implemented by SciBet.

To illustrate that SciBet is able to achieve high accuracy for self-projection within a dataset, 8 previously published datasets (Supplementary Table 3) were used to perform cross-validation tasks. SciBet achieved the highest accuracy compared to scmap^1^ and Seurat v3^2^ (Fig. 2b and Supplementary Fig. 2). To benchmark the performance including classification accuracy as well as efficiency, we generated 6 instances of training-test pairs using four human pancreas datasets generated by distinct sequencing techniques, and projected cells from one dataset onto annotated clusters from another dataset (Supplementary Table 4). For accuracy, SciBet showed a slight edge over Seurat v3 and scmap. For speed, SciBet out-performed others by orders of magnitude (Fig. 2c and Supplementary Fig. 3). We also tested whether SciBet could be used for cross-species classification, by projecting cells from three human pancreas datasets^6–8^ onto the *Tabula Muris* dataset^9^. Only genes with one-to-one orthologues cross-species were used for feature selection and cell identification (Online Methods). Most cells (∼92%) could be correctly mapped based on SciBet (Fig. 2d), demonstrating the utility of SciBet in cross-species cell identification.

Due to the incomplete nature of reference scRNA-seq data collection, cell types excluded from the reference dataset may be falsely predicted to be a known cell type. By applying a null dataset as background, which is generated by mixing all cell types together (Online Methods), SciBet controls the potential false positives while maintaining high prediction accuracy for cells with types covered by the reference dataset (positive cells). Using a recent melanoma dataset^10^ as an example, we showed that SciBet consistently provided the best performance by achieving low false positive rate as well as high accuracy for the positive cells (Fig. 2e). Although Seurat v3 also properly controlled false positives, most NK cells were incorrectly assigned to CD8 T cells or the “unassigned” group, leading to false negatives. Based on additional 16 previously published datasets (Supplementary Table 5), our evaluations demonstrated that SciBet correctly categorized >90% cells when the false positive rate (FPR) ranges from 0.001 to 0.05 (Fig. 2f). Both scmap and Seurat v3 had a much lower accuracies with the same FPR cutoffs, indicating the superiority of SciBet in prediction accuracy and false positive control (Fig. 2f and Supplementary Fig. 4).

We further collected 42 published human scRNA-seq datasets with full-length mRNA coverage and built an integrated dataset to include major human cell types that could be separated clearly by self-projection with SciBet (Fig. 2g). This dataset, analogous to Tabula Muris^9^, could serve as a plausible “mock” human cell atlas. We projected a recent human liver cell 10x genomics dataset^11^ to the integrated data by SciBet. Sankey diagram revealed that major cell types were correctly predicted, including hepatocytes, endothelial cells and Kupffer cells (specialized macrophages located in the liver). Besides, closely related cell types such as NK and T cells, as well as B and Plasma B cells, could also be precisely discriminated, suggesting the sensitivity and precision of SciBet for cross-platform prediction (Fig. 2h). Furthermore, each major cell type could be further classified into the minor label based on its corresponding dataset. Moreover, we provide a web server to upload custom data for such prediction. For large query dataset that may take a long time for data transmission, we also provide a lightweight standalone package for local construction of the web-based tools by a simple command.

Based on ∼100 well-annotated scRNA datasets from Single Cell Portal, Single Cell Expression Atlas, Gene Expression Omnibus and CellBlast^17^, we used SciBet to generate trained models for each dataset. The lightweight nature of trained models enabled the easy download together with the local SciBet package. For example, the size of a model with 100 cell types and 1000 signature genes is no more than 1 MB. We further built a JavaScript version of SciBet (http://scibet.cancer-pku.cn), which bypasses the process of file uploading to a remote server. This way, data files are read and processed locally and directly in the web browser, with only small-sized models transferred from the server to the browser, thus achieving unprecedented speed and convenience.

SciBet addresses an important need in the rapidly evolving field of single-cell transcriptomics, *i*.*e*., to accurately and rapidly capture main features of diverse datasets regardless of technical factors or batch effect. Based on multiple benchmarks, SciBet achieves high prediction accuracy, while keeping low false positive rate f or cells not represented previously. This advantage is achieved by considering not only the relative similarity to each cell type within the reference set, which is used by scmap^1^ and Seurat v3^2^ as well, but also the absolute similarity to the entire reference set and null dataset (Online Methods). Both E-test and SciBet utilizes the fundamental concept that each cell type is represented by the mean expression vector, as the computational clustering methods likens the experimental purification methods. Thus, both only carry out a small number of linear operations on the expression matrix, and the entire computational process is extremely efficient and can be thereby applied to very large-scale single cell studies emerging in coming years.

## ACKNOWLEDGMENTS

This project was supported by Beijing Advanced Innovation Centre for Genomics at Peking University, Key Technologies R&D Program (2016YFC0900100), National Natural Science Foundation of China (31530036 and 91742203). We thank the Computing Platform of the CLS (Peking University) for providing computing resource.

## AUTHOR CONTRIBUTIONS

Z.Z. conceived this study. C.L. developed the algorithm of E-test and SciBet; B.L. performed the benchmark testing. C.L., Z.L. and C.C developed the webserver. B.L. and B.K. developed the R package. C.L., B.L., X.R. and Z.Z. wrote the manuscript with all the authors’ inputs.

## COMPETING INTERESTS

No competing interests declared.

## ONLINE METHODS

### Data collection and processing

All scRNA-seq datasets and annotated cluster information used in this paper were obtained from their public accessions. For 26 datasets listed in Supplementary Table 1, we downloaded raw scRNA-seq data, and estimated the transcript-level abundance with kallisto^13^ and human genome reference hg19 (downloaded from UCSC).

### Parameterization of gene expression distribution of single cell data

We assume that the single cell expression level of a given gene follows the negative binomial distribution, which also arises as a continuous mixture of Poisson distributions with the gamma-distributed Poisson rate. Huang et al.^14^ models UMI count with this Poisson–Gamma mixture: Xgc∼Possion (λ_gc_*s_gc_), λ_gc_ ∼Gamma(α_gc_,β_gc_), where λ_gc_ represents the normalized true expression of gene g and cell c. We use CPM of each gene and cell as the moment estimation for the true expression level to eliminate the bias of library size. We generalize this representation form to full-length data, with TPM performing as the moment estimation for the true expression level after the normalization to eliminate the bias of gene length and library size. Thus, the normalized data follows the gamma distribution. For parameterizing α and β, we use one of the assumption that α keeps constant for all genes. Pseudo TPM or CPM 1 has been added to all element to the expression matrix.

### E-test for Measuring the difference of dispersion degree among cell groups

For a gamma distributed variable Y, the differential entropy can be calculated by the following formula: S(Y)= α-ln β+ln Γ(α)+(1-α) Ψ (α), where Ψ is the digamma function. The parameterβ can be estimated as α/E(Y), by maximum likelihood estimation. So the expression of S(Y) can be rewritten as S(Y)=lnE(Y)+g(α), where g(α)= α-ln α+ln Γ (α)+(1-α) Ψ (α), and performing as a constant according to our assumption that α is a constant.

For each gene, we calculate the statistic differential entropy (*S*) of its expression distribution across all cells, to reflect the degree of dispersion. As the distribution can be approximated by a negative binomial distribution, we can derive that the entropy statistic is approximately equal to the sum of the log mean expression (*E*) and a gene-independent constant (*S*-*E* formula, Methods). Given the expression levels of a gene across two or more different cell populations, a new statistic Δ*S* was designed to measure the total entropy reduction of all cell populations relative to a control cell population, mixed by all cells with group information removed. To highlight the differences among cell populations, we assume that there is no heterogeneity within each population, and hence *S* could be directly calculated by feeding its corresponding *E* into the *S*-*E* formula.

We calculate the sum of difference between each real cell groups and the null cell group, where there is no heterogeneity and the mean expression can be estimated as the average of the mean expression of all cell types. For datasets with m cell types, supervised entropy reduction of each gene could be calculated as:

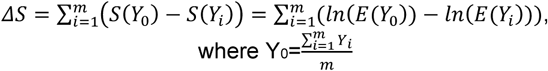

Under the null hypothesis that there is no difference among all cell populations, in other words each population is randomly sampled from the control population, Δ*S* would be zero-valued. A higher Δ*S* positive value would indicate higher differences among cell populations. Then genes with the top 1000 maximal Δ*S* were selected by default. This procedure is implemented in the DsGene function in SciBet. We observed that *ΔS*′ = log (*ΔS*) follows a normal distribution, so the significance of each gene can be evaluated with one-sided t-test.

### The multinomial assumption of expression levels across genes

We applied the multinomial distribution to model the process of mRNA generation of cell types. Assuming that mRNA molecules from the same gene are equivalent and that the generation for each mRNA molecule is independent, we can denote the probability of a newly generated mRNA molecule belonging to gene i as pi, and then each specific cell type can be represented by a probability vector p with pi as the elements. The vector p will serve as the parameters of a multinomial distribution to estimate the likelihood of a cell belonging to the given cell type.

### Estimation for the parameters of the multinomial distribution

Supposing there are m cells with identical cell type in the training dataset, and c_ij_ be the expression value of gene i ∈1,…,n in cell j∈1,…,m. Then the probability variable pi can be estimated by the mean vector across cells with normalization and Laplace Smoothing, which can be written as 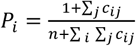. The smoothing accounts for genes not present in the expression matrix of training data which would cause zero probabilities and meaningless result for the following computations. This model has the least number of parameters theoretically with only one parameter per cell type per gene, so it can reduce the risk of over-fitting with similar goodness of fit. For example, when meeting a simulating drop-out events with a random portion of the matrix set to zero. The mean vector across cells will be in proportion to the origin one and the same after normalization. We use the log-transformed TPM or CPM to represent the true expression level.

### Bayesian decision on cell types for a testing cell

Supposing we have T cell types and each type has a probability vector p, the probability of a cell x of the test set with gene expressions x1,..,xn (n is the gene number) belonging to type y can be calculated by 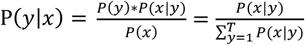, where 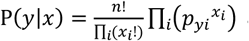 according to the multinomial assumption and equals 1/T for each given y because we do not consider about the prior probability of class. We identified type y with the largest probability value of cell x as the predicted cell label, which can be written as 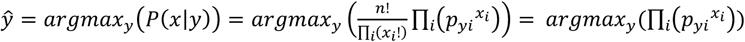. It is worth noting that if a dropout event occurs for one gene in the testing cell, the value zero of xi will lead to *p*_*i*_ ^0^=1 and thus does not affect other genes.

### The connection between the multinomial distribution and the cross Entropy

The abovementioned 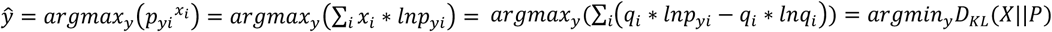, where *q*_*i*_ = *x*_*i*_/Σ_*i*_ *x*_*i*_. It indicates that we annotate the test cell to such the cell type that its mean expression vector of all marker genes has the minimal Kullback–Leibler divergence to the test cell.

### False positive control

To control potential false positive predictions, we first identified common marker genes (n=500) exclusively expressed in reference cells using E-test, with a null dataset (an integrated dataset using 35 studies listed in Supplementary Table 1) served as the background. Then we calculate the occurrence probability of each marker gene in reference set and null set respectively as (3). In this way, the classification score of each query cell could be defined as:

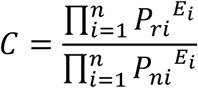

where *E*_*i*_ is the expression value of gene *i* in cell *x*, and *P*_*ri*_ and *P*_*ni*_ are occurrence probability values of gene *i* in reference and null set, respectively.

For false positive control benchmarking experiments (Fig. 2f and Supplementary Fig. 4), we assigned “unassigned” to the cells with lowest classification scores with a series of cutoffs from 0.001 to 0.5.

### Seurat v3, scmap, SVM and RF

SVM^15^ (with a linear kernel) and RF^16^ (with 100 trees) from the python module sklearn were used in our analysis. For scmap^1^, we performed the classification using the function scmapCluster and set the similarity threshold to 0, with n_features specified as 500. For Seurat v3^2^, we identified anchors and classified cells using the FindTransferAnchors function and TransferData functions, respectively, with default parameters.

### Cross species classification

We used the HomoloGene databases provided by NCBI (Build 68) to identify homologous genes between human and mouse, and kept only genes that have one-to-one correspondence, which works as a dictionary. After getting the intersection gene set between the training dataset (Tabula Muris) and the dictionary, the gene names were then converted to human gene names to get a test-set-compatible training set for classification.

### Implementation and webserver

All the functions mentioned above were implemented in the software package SciBet written with R, which be download at http://scibet.cancer-pku.cn. An online version of SciBet is available at http://scibet.cancer-pku.cn, which is based on JavaScript.

## Main Figures

**Supplementary Figure 1. E-test performance**. E-test performance on n = 14 cases (Supplementary Table 1) is measured by percentage of correctly mapped cells, with **(a)** SVM, **(b)** RF and **(c)** SciBet, corresponding to Fig. 1b.

**Supplementary Figure 2. SciBet performance on self-projections**. Classification accuracy of self-projection tasks, estimated by SciBet, Seurat v3 and scmap for n = 8 datasets listed in Supplementary Table 3, corresponding to Fig. 2b.

**Supplementary Figure 3. SciBet performance on cross platforms**. Speed and classification accuracy of cross-platform experiments estimated by SciBet, Seurat v3 and scmap with 6 instances (Supplementary Table 4), corresponding to Fig. 2c.

**Supplementary Figure 4. SciBet performance on false positive contral**. False positive evaluation with cell types not present in reference as negative control, estimated by SciBet, Seurat v3 and scmap with n = 8 cases (Supplementary Table 5).

## Notes

#### Summary of Updates

Web server updated (with JavaScript implementation of SciBet); Improvement of SciBet performance; Figure 2 revised; Author affiliations updated; Supplemental files updated.

