## Supplementary figures and images for "SciBet: a portable and fast single cell type identifier"

### Supplementary figure 1

Supplementary Figure 1

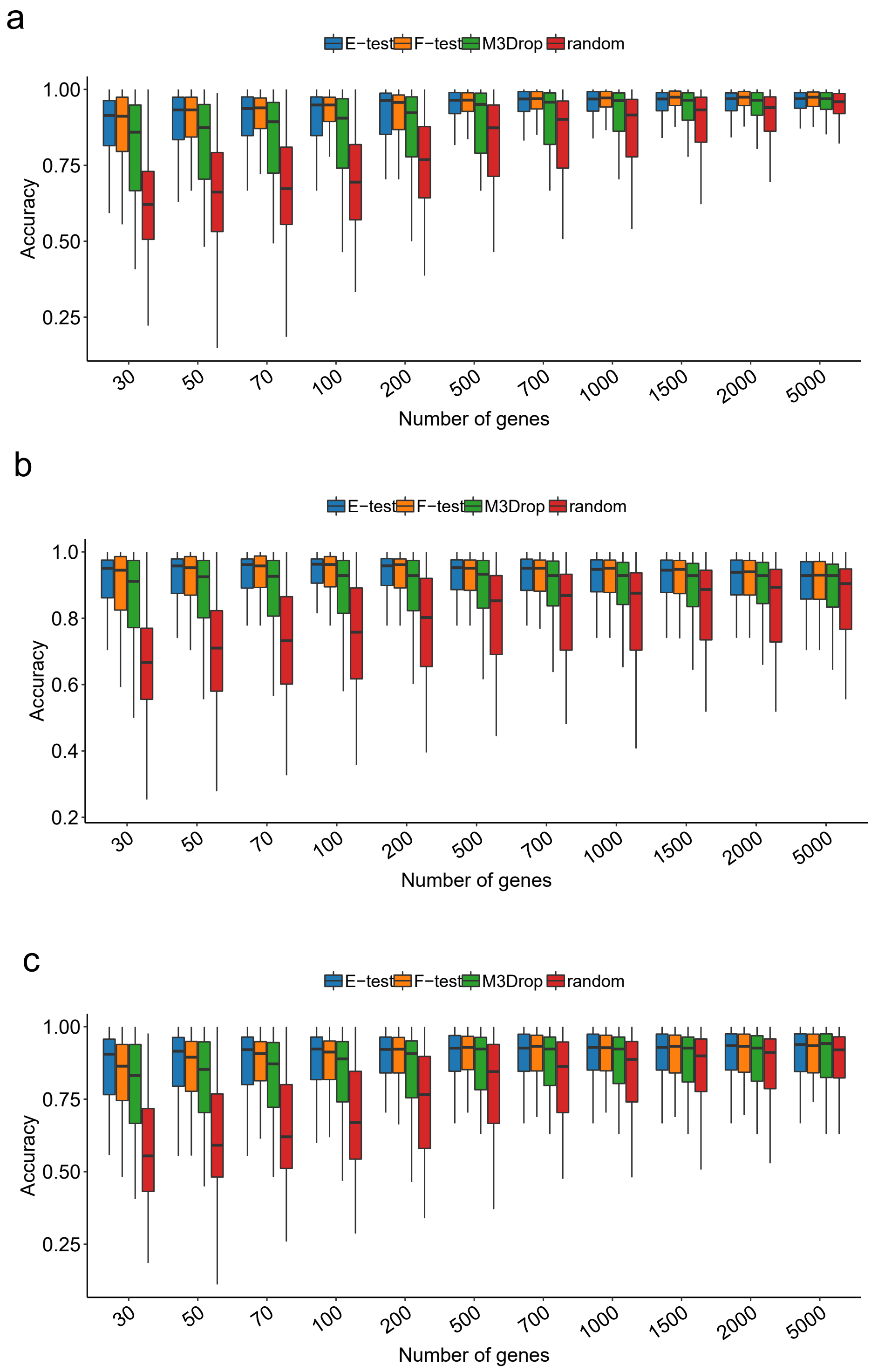

### Supplementary figure 2

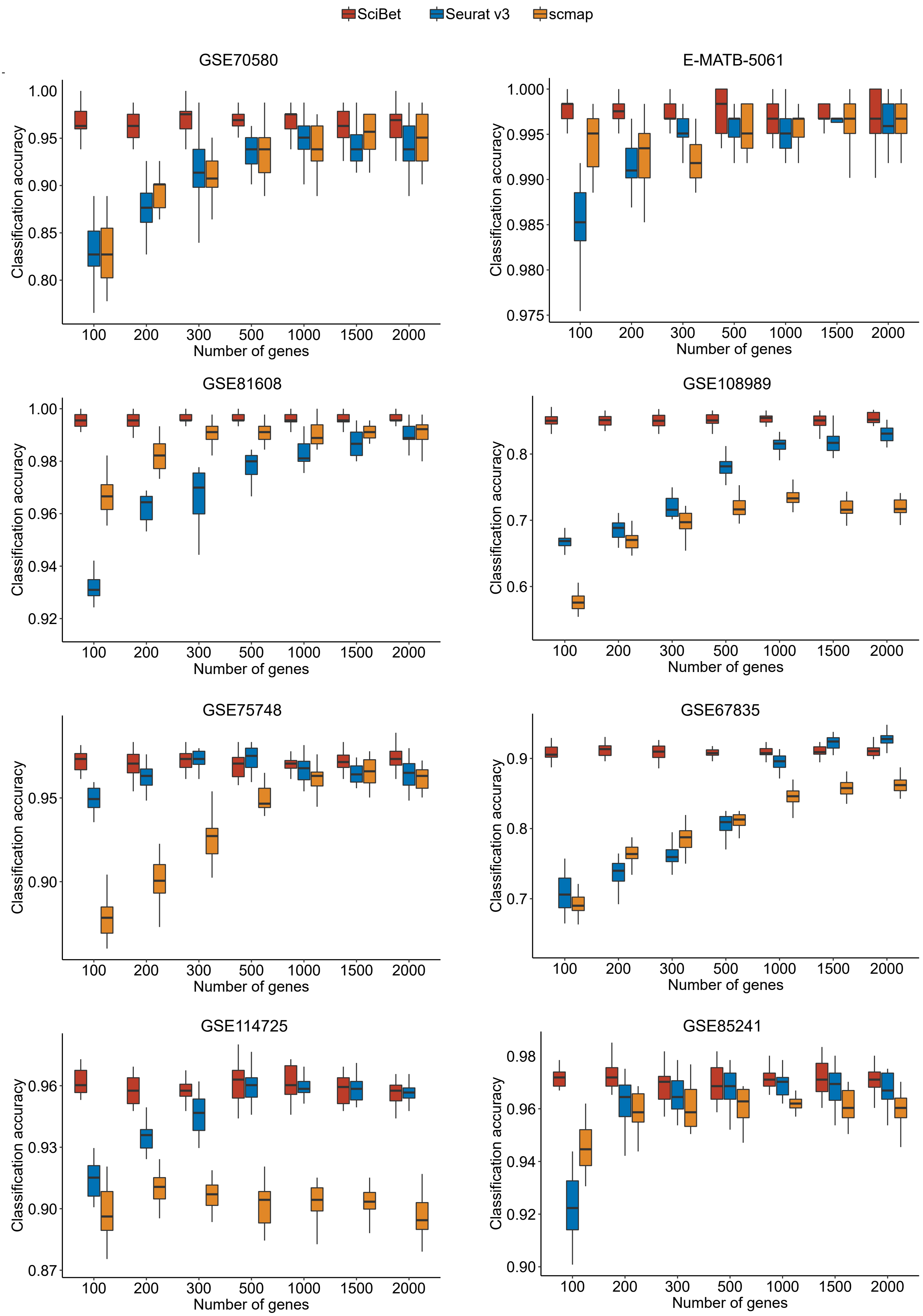

### Supplementary figure 3

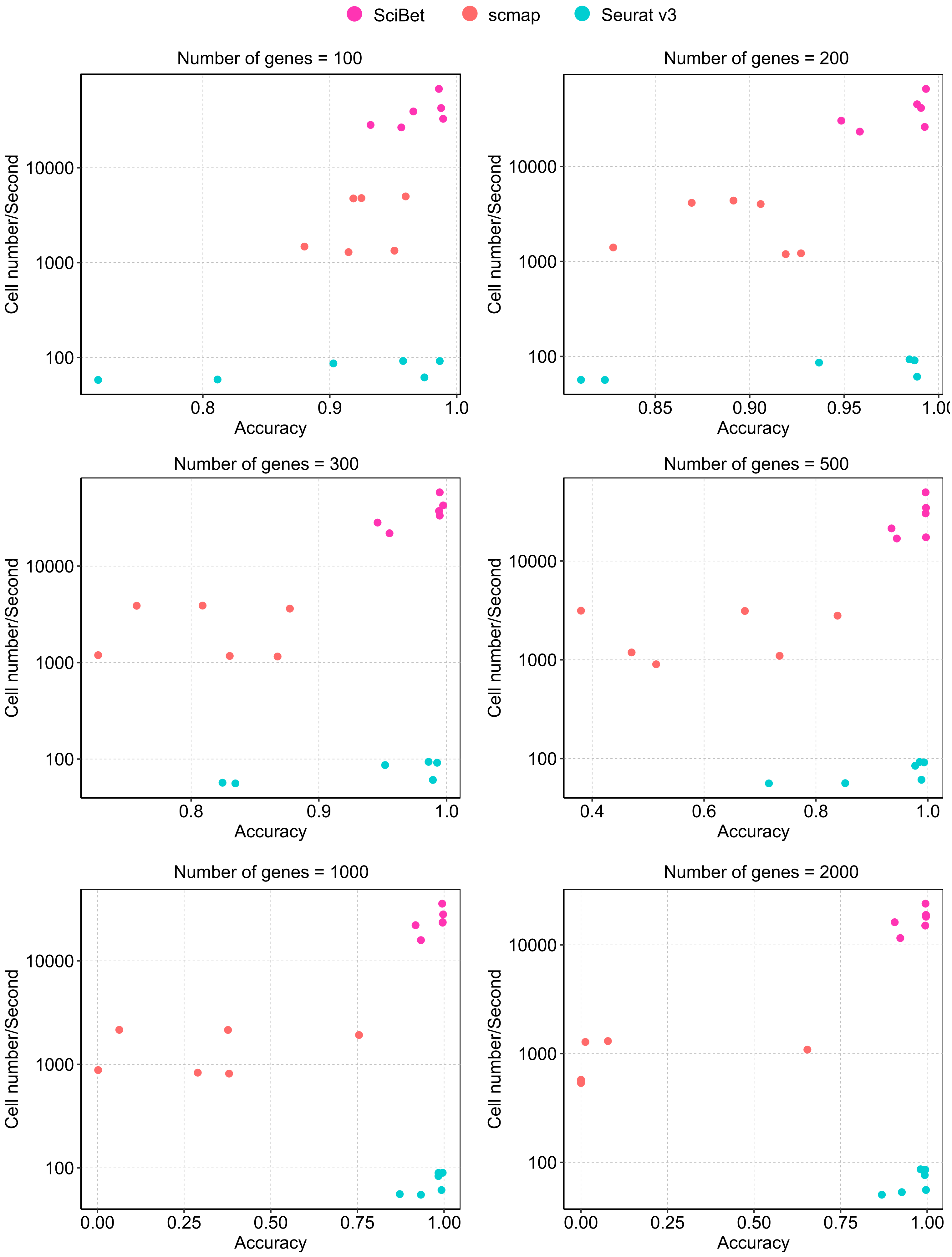

### Supplementary figure 4

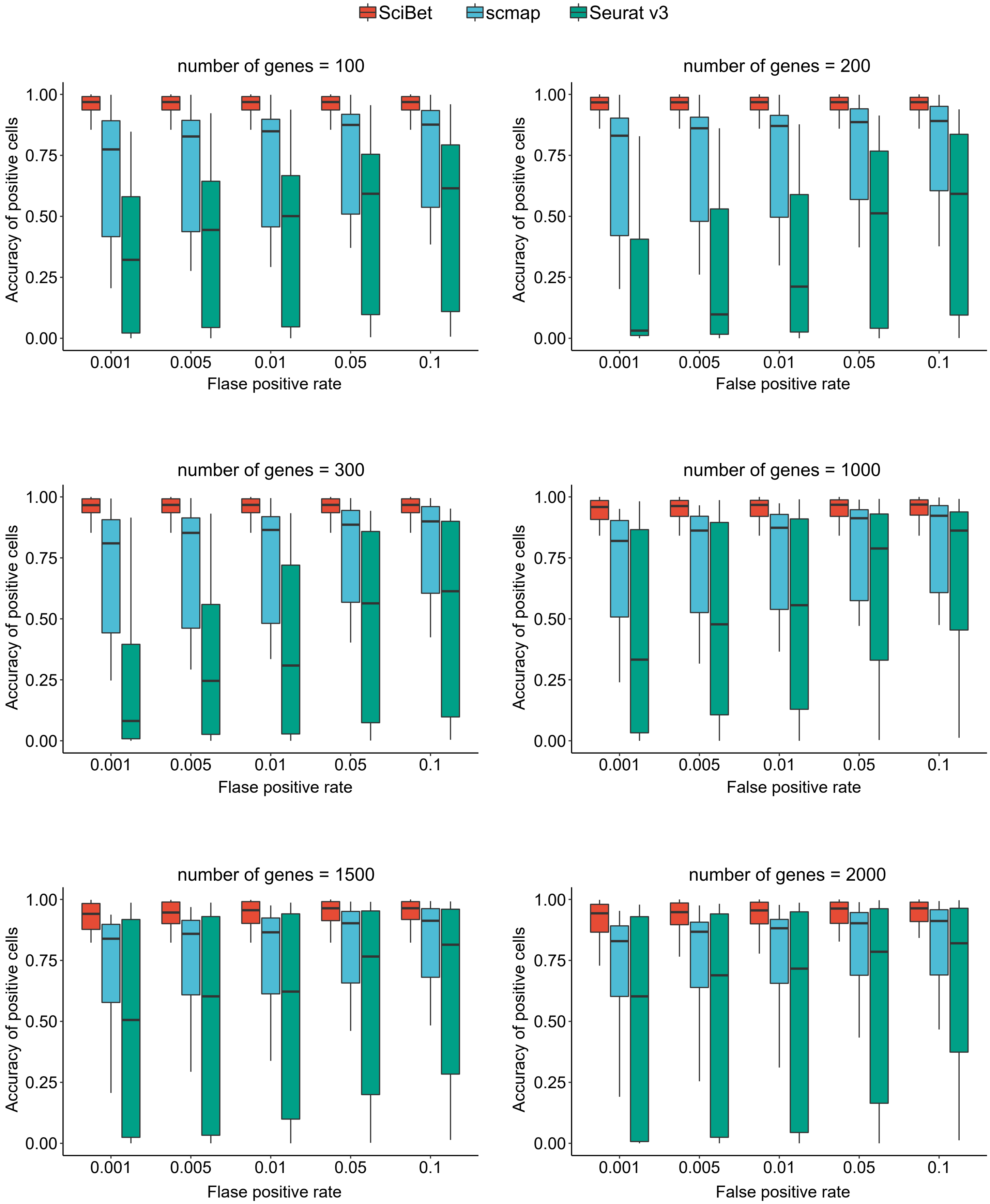
